# Reduced antibody cross-reactivity following infection with B.1.1.7 than with parental SARS-CoV-2 strains

**DOI:** 10.1101/2021.03.01.433314

**Authors:** Nikhil Faulkner, Kevin W. Ng, Mary Wu, Ruth Harvey, Marios Margaritis, Stavroula Paraskevopoulou, Catherine F. Houlihan, Saira Hussain, Maria Greco, William Bolland, Scott Warchal, Judith Heaney, Hannah Rickman, Moira J. Spyer, Daniel Frampton, Matthew Byott, Tulio de Oliveira, Alex Sigal, Svend Kjaer, Charles Swanton, Sonia Gandhi, Rupert Beale, Steve J. Gamblin, Crick COVID-19 Consortium, John McCauley, Rodney Daniels, Michael Howell, David L.V. Bauer, Eleni Nastouli, SAFER Investigators, George Kassiotis

## Abstract

We examined the immunogenicity of severe acute respiratory syndrome coronavirus 2 (SARS-CoV-2) variant B.1.1.7 that arose in the United Kingdom and spread globally. Antibodies elicited by B.1.1.7 infection exhibited significantly reduced recognition and neutralisation of parental strains or of the South Africa B.1.351 variant, than of the infecting variant. The drop in cross-reactivity was more pronounced following B.1.1.7 than parental strain infection, indicating asymmetric heterotypic immunity induced by SARS-CoV-2 variants.

## Main

Mutations in SARS-CoV-2 variants that arose in the United Kingdom (UK) (B.1.1.7) or in South Africa (B.1.351) reduce recognition by antibodies elicited by natural infection with the parental reference (Wuhan) strain and the subsequent D614G variant^1–10^. Such reduction in cross-reactivity also impinges the effectiveness of current vaccines based on the Wuhan strain^4–10^, prompting consideration of alternative vaccines based on the new variants. However, the immunogenicity of the latter or, indeed, the degree of heterotypic immunity the new variants may afford remains unknown.

The B.1.1.7 variant is thought to have first emerged in the UK in September 2020 and has since been detected in over 50 countries^11^. To examine the antibody response to B.1.1.7, we collected sera from 29 patients, admitted to University London College Hospital (UCLH) for unrelated reasons (Table S1), who had confirmed B.1.1.7 infection. The majority (23/29) of these patients displayed relatively mild COVID-19 symptoms and a smaller number (6/29) remained COVID-19-asymptomatic. As antibody titres may depend on the severity of SARS-CoV-2 infection, as well as on time since infection, we compared B.1.1.7 sera with sera collected during the first wave of D614G variant spread in London from hospitalised COVID-19 patients^12^ (n=20) and asymptomatic SARS-CoV-2-infected health care workers^13^ (n=17) who were additionally sampled two months later.

IgG, IgM and IgA antibodies to the spikes of the Wuhan strain or of variants D614G, B.1.1.7 or B.1.351, expressed on HEK293T cells, were detected by a flow cytometry-based method (Fig. S1)^12^. Titres of antibodies that bound the parental D614G spike largely correlated with those that bound the B.1.1.7 or B.1.351 spikes (Fig. 1a-c), consistent with the high degree of similarity. Similar correlations were observed for all three Ig classes also between the Wuhan strain and the three variant spikes and between the B.1.1.7 and B.1.351 spikes (Fig. S2-S5).

**Figure 1.**
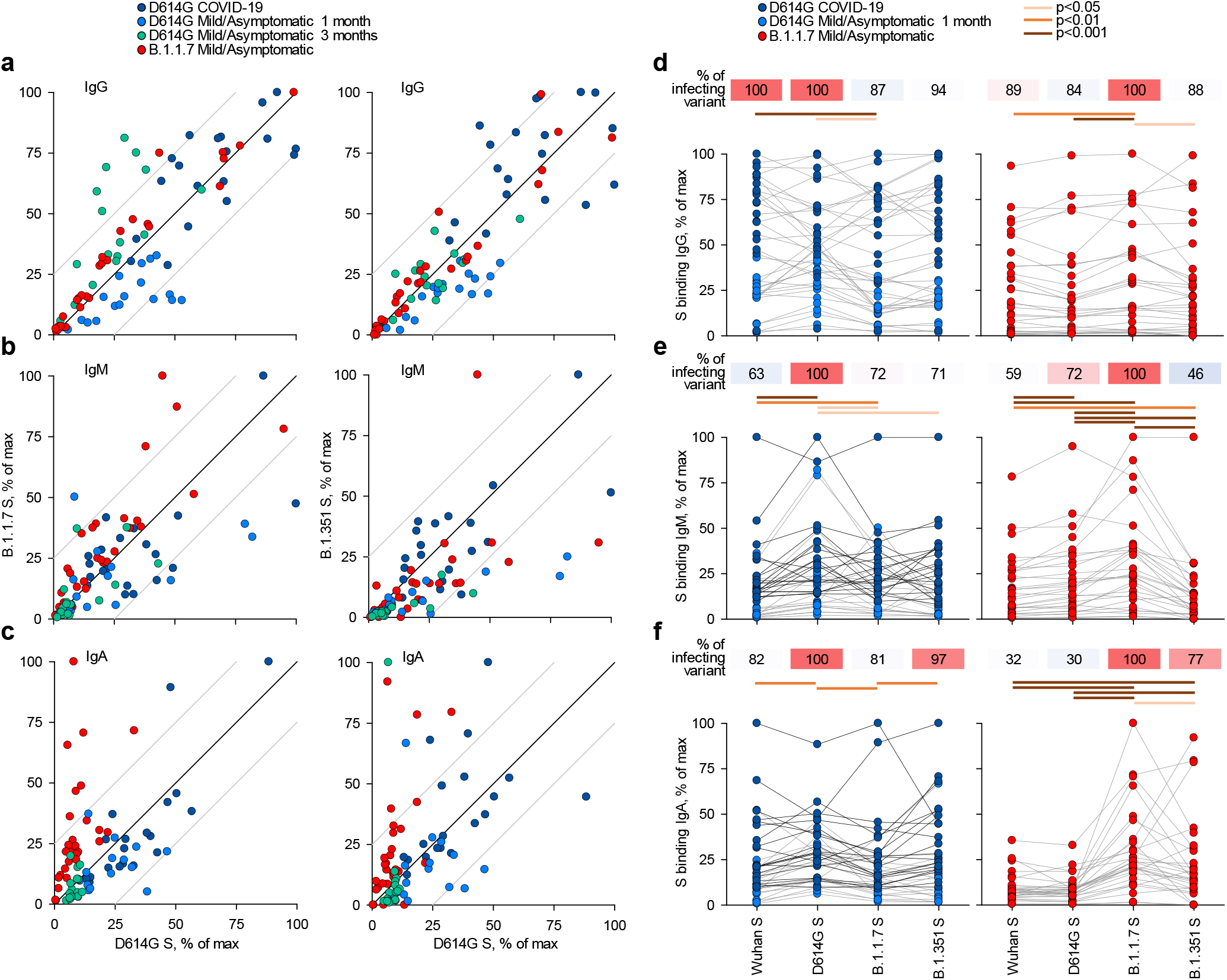
Recognition of distinct SARS-CoV-2 spike glycoproteins by antibodies in D614G and B.1.1.7 sera. **a-c**, Correlation of IgG (a), IgM (b) and IgA (c) antibody levels to D614G and B.1.1.7 or B.1.351 spikes in the indicated groups of donors infected either with the D614G or B.1.1.7 strains. Each symbol represents an individual sample and levels are expressed as a percentage of the positive control. Black lines denote complete correlation and grey lines a 25% change in either direction. **d-f**, Comparison of IgG (d), IgM (e) and IgA (f) antibody levels to the indicated spikes in groups of donors acutely infected either with the D614G or B.1.1.7 strains. Connected symbols represents individual donors. Numbers above the plots denote the average binding to each spike, expressed as a percentage of binding to the infecting spike.

Comparison of sera from acute D614G and B.1.1.7 infections revealed stronger recognition of the infecting variant than of other variants. Although B.1.1.7 sera were collected on average earlier than D614G sera (Table S1), titres of antibodies that bound the homotypic spike or neutralised the homotypic virus, as well as the relation between these two properties, were similar in D614G and B.1.1.7 sera (Fig. S6), suggesting comparable immunogenicity of the two variants. Recognition of heterotypic spikes was reduced by a small, but statistically significant degree for both D614G and B.1.1.7 sera and for all three Ig classes (Fig. 1d-f). IgM or IgA antibodies in both D614G and B.1.1.7 sera were less cross-reactive than IgG antibodies (Fig. 1d-f). The direction of cross-reactivity was disproportionally affected for some combinations, with IgA antibodies in D614G sera retaining on average 81% of recognition of the B.1.1.7 spike and IgA antibodies in B.1.1.7 sera retaining on average 30% of recognition of the D614G spike (Fig. 1f). Similarly, recognition of the B.1.351 spike by IgM antibodies was retained, on average, to 71% in D614G sera and to 46% in B.1.1.7 sera (Fig. 1f). Measurable reduction in polyclonal antibody binding to heterotypic spikes was unexpected, given >98% amino acid identity between them. However, reduction in serum antibody binding has also been observed for the receptor binding domain of the B.1.351 spike^4^. Together, these findings suggested that either the limited number of mutated epitopes were targeted by a substantial fraction of the response^7–10^ or allosteric effects or conformational changes affecting a larger fraction of polyclonal antibodies.

To examine a functional consequence of reduced antibody recognition, we measured the half maximal inhibitory concentration (IC_50_) of D614G and B.1.1.7 sera using *in vitro* neutralisation of authentic Wuhan or B.1.1.7 and B.1.351 viral isolates (Fig. 2a-b). Titres of neutralising antibodies correlated most closely with levels of IgG biding antibodies for each variant (Fig. S5). Neutralisation of B.1.1.7 by D614G sera was largely preserved at levels similar to neutralisation of the parental Wuhan strain (fold change −1.3; range 3.0 to −3.8, p=0.183) (Fig. 2b), consistent with other recent reports, where authentic virus neutralisation was tested^1,7–9,14^. Moreover, the ability of D614G sera to neutralise B.1.1.7 was preserved over time (Fig. S7). Indeed, whilst binding antibody titres were significantly reduced for all three Ig classes in D614G sera in the two months of follow-up, neutralising antibody titres remained comparable for the Wuhan and B.1.1.7 strains and were undetectable at both time-points for the B.1.351 strain (Fig. S7). Thus, D614G infection appeared to induce substantial and lasting cross-neutralisation of the B.1.1.7 variant. However, the reverse was not true. Neutralisation of the parental Wuhan strain by B.1.1.7 sera was significantly reduced, compared to neutralisation of the infecting B.1.1.7 variant (fold change −3.4; range −1.20 to −10.6, p<0.001) (Fig. 2b), and the difference in cross-neutralisation drop was also significant (p<0.001). Both D614G and B.1.1.7 sera displayed significantly reduced neutralisation of the B.1.351 variant with a fold change of −8.2 (range −1.7 to −33.5) and −7.7 (range −3.4 to −17.9), respectively (Fig. 2b).

**Figure 2.**
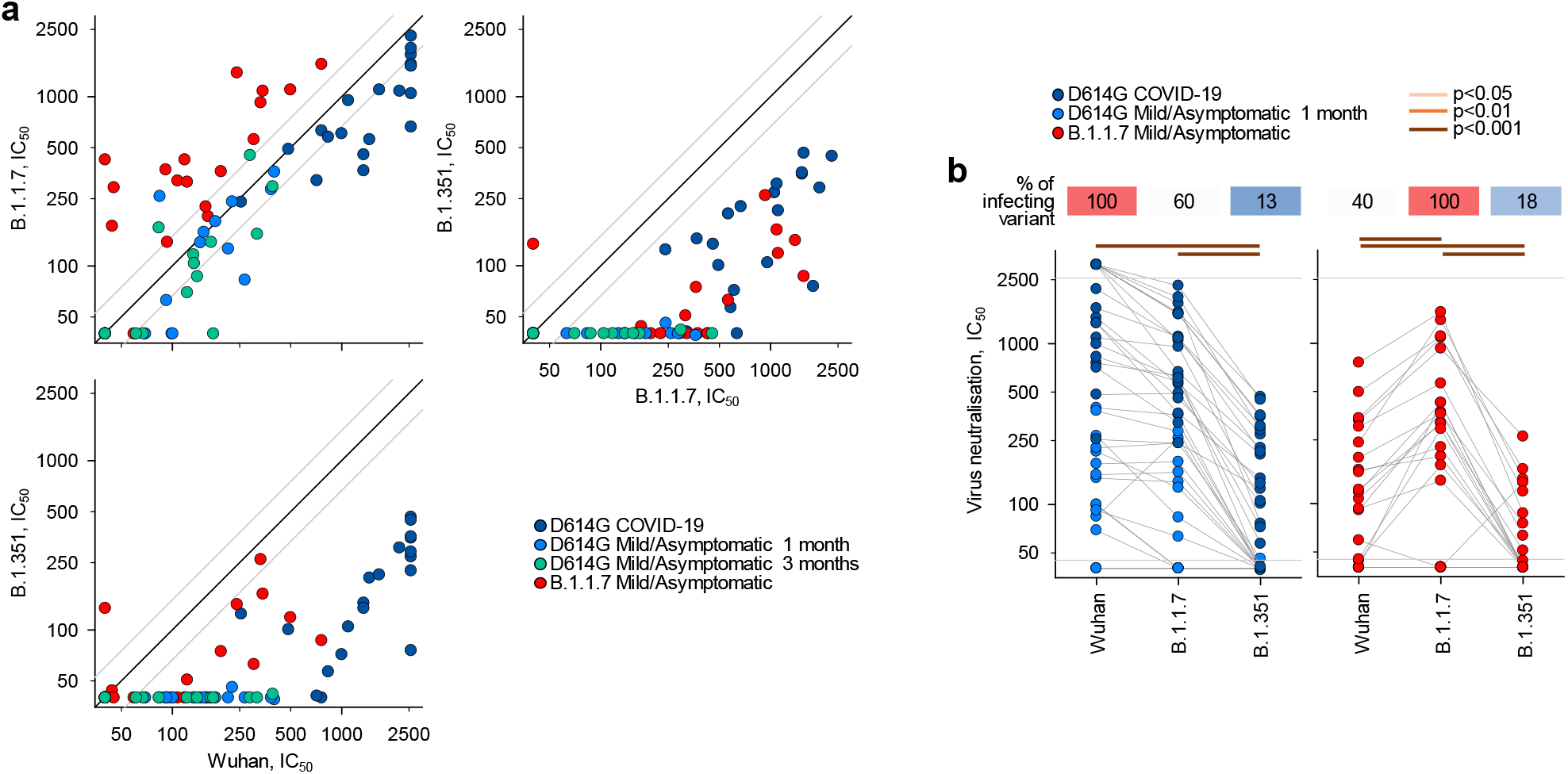
Neutralisation of distinct SARS-CoV-2 strains by antibodies in D614G and B.1.1.7 sera. **a**, Correlation of neutralising antibody levels (IC_50_) against the Wuhan, B.1.1.7 or B.1.351 strains in the indicated groups of donors infected either with the D614G or B.1.1.7 strains. Each symbol represents an individual sample. Black lines denote complete correlation and grey lines a 50% (2-fold) change in either direction. **b**, Comparison of neutralising antibody levels (IC_50_) to the indicated SARS-CoV-2 strains in groups of donors acutely infected either with the D614G or B.1.1.7 strains. Connected symbols represents individual donors. Numbers above the plots denote the average IC_50_ against each strain, expressed as a percentage of IC_50_ against the infecting strain. Grey horizontal lines denote the lower and upper limit of detection.

Together, these results argue that natural infection with each SARS-CoV-2 strain induces antibodies that recognise the infecting strain most strongly, with variable degrees of cross-recognition of the other strains. Importantly, antibodies induced by B.1.1.7 infection were less cross-reactive with other dominant SARS-CoV-2 strains than those induced by the parental strain. Similar findings were recently obtained independently with a small number of B.1.1.7 convalescent sera^14^. This unidirectional pattern of cross-reactivity argues that emergence of B.1.1.7 is unlikely to have been driven by antibody escape.

A recent comparison of sera from infection with B.1.351 or the parental strain B.1.1.117 in South Africa, also observed stronger neutralisation of the infecting strain^3^. In contrast to B.1.1.7 infection, however, B.1.351 infection induced substantial cross-neutralisation of the parental strain, whereas parental strain B.1.1.117 infection induced significantly lower B.1.351 neutralisation^3^. Therefore, heterotypic immunity in the case of B.1.351 and the parental strain B.1.1.117 was also asymmetrical, but reversed.

Although a quantifiable correlation between *in vitro* neutralisation of infectious SARS-CoV-2 and *in vivo* protection from SARS-CoV-2 infection or severe COVID-19 remains to be defined^15^, the reduced neutralisation of other SARS-CoV-2 strains by B.1.1.7 sera would suggests that the recent wave of global B.1.1.7 infections may not completely protect against re-infection with other SARS-CoV-2 strains. The degree of heterotypic immunity should be an important consideration in the choice of spike variants as vaccine candidates, with B.1.1.7 spike demonstrating lower potential than other variants. The antigenic variation associated with SARS-CoV-2 evolution may instead necessitate the use of multivalent vaccines.

## Methods

### Donor and patient samples and clinical data

Serum or plasma samples from D614G infection were obtained from University College London Hospitals (UCLH) (REC ref: 20/HRA/2505) COVID-19 patients (n=20, acute D614G infection, COVID-19 patients) as previously described^12^, or from UCLH health care workers (n=17, acute D614G infection, mild/asymptomatic), as previously described^13^ (Table S1). These samples were collected between March 2020 and June 2020. Serum or plasma samples from B.1.1.7 infection were obtained from patients (n=29, acute B.1.1.7 infection, mild/asymptomatic) admitted to UCLH (REC ref: 20/HRA/2505) for unrelated reasons, between December 2020 and January 2021, who then tested positive for SARS-CoV-2 infection by RT-qPCR, as part of routine testing (Table S1). Infection with B.1.1.7 was confirmed by sequencing of viral RNA, covered from nasopharyngeal swabs. A majority of these patients (n=23) subsequently developed mild COVID-19 symptoms and 6 remained asymptomatic. All serum or plasma samples were heat-treated at 56°C for 30 min prior to testing.

### Diagnosis of SARS-CoV-2 infection by RT-qPCR and next generation sequencing

SARS-CoV-2 nucleic acids were detected in nasopharyngeal swabs from hospitalised patients by a diagnostic RT-qPCR assay using custom primers and probes^16^, Assays were run by Health Services Laboratories (HSL), London, UK. Diagnostic RT-qPCR assays for SARS-CoV-2 infection in health care workers was run at the Francis Crick Institute, as previously described^17^. SARS-CoV-2 RNA-positive samples (RNA amplified by Aptima Hologic) were subjected to real-time whole-genome sequencing at the UCLH Advanced Pathogen Diagnostics Unit. RNA was extracted from nasopharyngeal swab samples on the QiaSymphony platform using the Virus Pathogen Mini Kit (Qiagen). Libraries were prepared using the Illumina DNA Flex library preparation kit and sequenced on an Illumina MiSeq (V2) using the ARTIC protocol for targeted amplification (primer set V3). Genomes were assembled using an in-house pipeline^18^ and aligned to a selection of publicly available SARS-CoV-2 genomes^19^ using the MAFFT alignment software^20^. Phylogenetic trees were generated from multiple sequence alignments using IQ-TREE^21^ and FigTree (http://tree.bio.ed.ac.uk/software/figtree), with lineages assigned (including B.1.1.7 calls) using pangolin (http://github.com/cov-lineages/pangolin), and confirmed by manual inspection of alignments.

### Cells lines and plasmids

HEK293T cells were obtained from the Cell Services facility at The Francis Crick Institute, verified as mycoplasma-free and validated by DNA fingerprinting. Vero E6 and Vero V1 cells were kindly provided by Dr Björn Meyer, Institut Pasteur, Paris, France, and Professor Steve Goodbourn, St. George’s, University of London, London, UK, respectively. Cells were grown in Iscove’s Modified Dulbecco’s Medium (Sigma Aldrich) supplemented with 5% fetal bovine serum (Thermo Fisher Scientific), L-glutamine (2 mM, Thermo Fisher Scientific), penicillin (100 U/ml, Thermo Fisher Scientific), and streptomycin (0.1 mg/ml, Thermo Fisher Scientific). For SARS-CoV-2 spike expression, HEK293T cells were transfected with an expression vector (pcDNA3) carrying a codon-optimized gene encoding the wild-type SARS-CoV-2 reference spike (referred to here as Wuhan spike, UniProt ID: P0DTC2) or a variant carrying the D614G mutation and a deletion of the last 19 amino acids of the cytoplasmic tail (referred to here as D614G spike) (both kindly provided by Massimo Pizzato, University of Trento, Italy). Similarly, HEK293T cells were transfected with expression plasmids (pcDNA3) encoding the B.1.1.7 spike variant (D614G, Δ69-70, Δ144, N501Y, A570D, P681H, T716I, S982A and D1118H) or the B.1.351 spike variant (D614G, L18F, D80A, D215G, Δ242-244, R246I, K417N, E484K, N501Y, A701V) (both synthesised and cloned by GenScript). All transfections were carried out using GeneJuice (EMD Millipore) and transfection efficiency was between 20% and 54% in separate experiments.

### SARS-CoV-2 isolates

The SARS-CoV-2 reference isolate (referred to as the Wuhan strain) was the hCoV-19/England/02/2020, obtained from the Respiratory Virus Unit, Public Health England, UK, (GISAID EpiCov™ accession EPI_ISL_407073). The B.1.1.7 isolate was the hCoV-19/England/204690005/2020, which carries the D614G, Δ69-70, Δ144, N501Y, A570D, P681H, T716I, S982A and D1118H mutations^14^, obtained from Public Health England (PHE), UK, through Prof. Wendy Barclay, Imperial College London, London, UK. The B.1.351 virus isolate was the 501Y.V2.HV001, which carries the D614G, L18F, D80A, D215G, Δ242-244, K417N, E484K, N501Y, A701V mutations^3^. However, sequencing of viral genomes isolated following further passage in Vero V1 cells identified the Q677H and R682W mutations at the furin cleavage site, in approximately 50% of the genomes. All viral isolates were propagated in Vero V1 cells.

### Flow cytometric detection of antibodies to spike and envelope glycoproteins

HEK293T cells were transfected to express the different SARS-CoV-2 spike variants. Two days after transfection, cells were trypsinized and transferred into V-bottom 96-well plates (20,000 cells/well). Cells were incubated with sera (diluted 1:50 in PBS) for 30 min, washed with FACS buffer (PBS, 5% BSA, 0.05% sodium azide), and stained with BV421 anti-IgG (clone HP6017, Biolegend), APC anti-IgM (clone MHM-88, Biolegend) and PE anti-IgA (clone IS11-8E10, Miltenyi Biotech) for 30 min (all antibodies diluted 1:200 in FACS buffer). Cells were washed with FACS buffer and fixed for 20 min in CellFIX buffer (BD Bioscience). Samples were run on a Ze5 analyzer (Bio-Rad) running Bio-Rad Everest software v2.4 or an LSR Fortessa with a high-throughput sampler (BD Biosciences) running BD FACSDiva software v8.0, and analyzed using FlowJo v10 (Tree Star Inc.) analysis software, as previously described^12^. All runs included 3 positive control samples, which were used for normalisation of mean fluorescence intensity (MFI) values. To this end, the MFI of the positively stained cells in each sample was expressed as a percentage of the MFI of the positive control on the same 96-well plate.

### SARS-CoV-2 neutralisation assay

SARS-CoV-2 variant neutralisation was tested using an in-house developed method (Fig. S8). Heat-inactivated serum samples in QR coded vials (FluidX/Brooks) were assembled into 96-well racks along with foetal calf serum-containing vials as negative controls and SARS-CoV-2 spike RBD-binding nanobody (produced in-house) vials as positive controls. A Viaflo automatic pipettor fitted with a 96-channel head (Integra) was used to transfer serum samples into V-bottom 96-well plates (Thermo 249946) prefilled with Dulbecco’s Modified Eagle Medium (DMEM) to achieve a 1:10 dilution. The Viaflo was then used to serially dilute from the first dilution plate into 3 further plates at 1:4 to achieve 1:40, 1:160, and 1:640. Next, the diluted serum plates were stamped into duplicate 384-well imaging plates (Greiner 781091) pre-seeded the day before with 3,000 Vero E6 cells per well, with each of the 4 dilutions into a different quadrant of the final assay plates to achieve a final working dilution of samples at 1:40, 1:160, 1:640, and 1:2560. Assay plates were then transferred to containment level 3 (CL3) where cells were infected with the indicated SARS-CoV-2 viral strain, by adding a pre-determined dilution of the virus prep using a Viaflo fitted with a 384 head with tips for the no-virus wells removed. Plates were incubated for 24 hours at 37°C, 5% CO_2_ and then fixed by adding a concentrated formaldehyde solution to achieve a final concentration of 4%. Assay plates were then transferred out of CL3 and fixing solution washed off, cells blocked and permeabilised with a 3% BSA/0.2% Triton- X100/PBS solution, and finally immunostained with DAPI and a 488-conjugated anti-nucleoprotein monoclonal antibody (produced in-house). Automated imaging was carried out using an Opera Phenix (Perkin Elmer) with a 5x lens and the ratio of infected area (488-positive region) to cell area (DAPI-positive region) per well calculated by the Phenix-associated software Harmony. A custom automated script runs plate normalisation by background subtracting the median of the no-virus wells and then dividing by the median of the virus-only wells before using a 3-parameter dose-response model for curve fitting and identification of the dilution which achieves 50% neutralisation for that particular serum sample (IC_50_).

### Statistical analyses

Data were analysed and plotted in SigmaPlot v14.0 (Systat Software). Parametric comparisons of normally-distributed values that satisfied the variance criteria were made by paired or unpaired Student’s t-tests or One Way Analysis of variance (ANOVA) tests. Data that did not pass the variance test were compared with Wilcoxon Signed Rank Tests.

## Acknowledgements

We are grateful for assistance from the Flow Cytometry and Cell Services facilities at the Francis Crick Institute and to Mr Michael Bennet and Mr Simon Caidan for training and support in the high-containment laboratory. We wish to thank the Public Health England (PHE) Virology Consortium and PHE field staff, the ATACCC (Assessment of Transmission and Contagiousness of COVID-19 in Contacts) investigators, the G2P-UK (Genotype to Phenotype-UK) National Virology Consortium, and Prof Wendy Barclay, Imperial College London, London, UK, for the B.1.1.7 viral isolate. This work was supported by the Francis Crick Institute, which receives its core funding from Cancer Research UK, the UK Medical Research Council, and the Wellcome Trust. The funders had no role in study design, data collection and analysis, decision to publish, or preparation of the manuscript.

---

**Table S1.** Donor and patient characteristics.

| **Donor/patient group** | **Number** | **Age, years**<br>**(range)** | **Male gender**<br>**(%)** | **Days post infection<sup>2</sup>,**<br>**median (range)** |
| --- | --- | --- | --- | --- |
| Acute D614G infection,<br>mild/asymptomatic<sup>1</sup> | 17 | 32<br>(26-66) | 7<br>(41) | 28<br>(6-35) |
| Acute D614G infection,<br>COVID-19 patients | 20 | 61<br>(25-73) | 14<br>(70) | 25<br>(14-43) |
| Acute B.1.1.7 infection,<br>mild/asymptomatic | 29 | 57<br>(20-99) | 18<br>(62) | 11<br>(4-46) |

<sup>1</sup>Seroconversion and/or first positive RT-qPCR in the first month of enrolment (month 1). The same group of participants were sampled again 2 months later (month 3).
<sup>2</sup>Days post symptom onset for symptomatic cases and days post first positive RT-qPCR test for asymptomatic cases.

**Figure S1.**
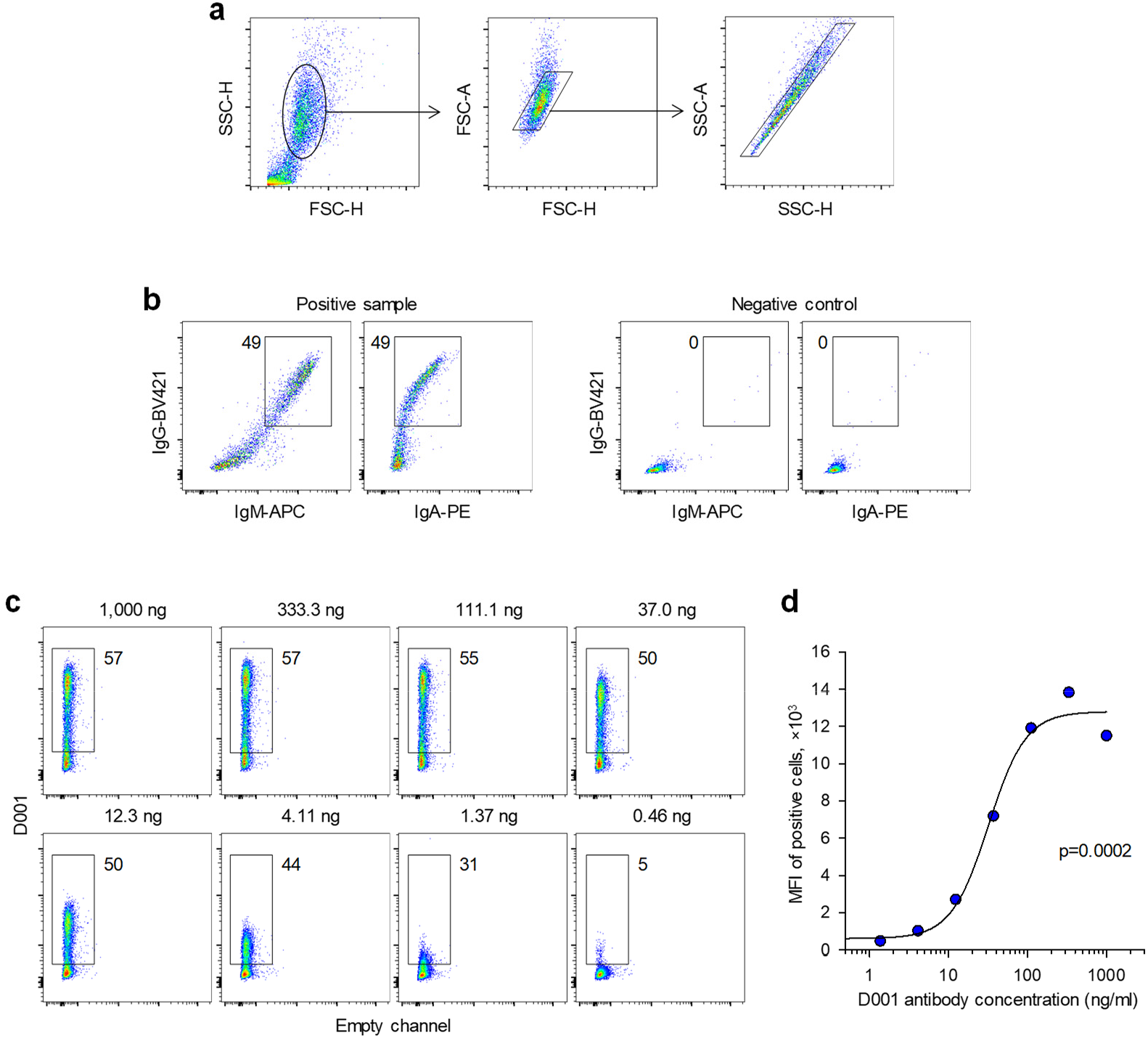
Flow cytometric detection of spike-binding antibodies. HEK293T cells were transfected with expression plasmids encoding each SARS-CoV-2 variant spike and were used for flow cytometric analysis two days later. **a**, Gating of HEK293T cells and of single cells in these mixed cell suspensions. **b**, Example of IgG, IgM and IgA staining in a positive sample and a negative control. Numbers within the plots denote the percentage of positive cells. **c**, Staining of HEK293T cells transfected to express the Wuhan spike, with titrated amounts of the S2-specific D001 monoclonal antibody. Numbers above the plots denote the final D001 antibody concentration. **d**, Median fluorescence intensity (MFI) of stained cells in c, according to the D001 antibody concentration.

**Figure S2.**
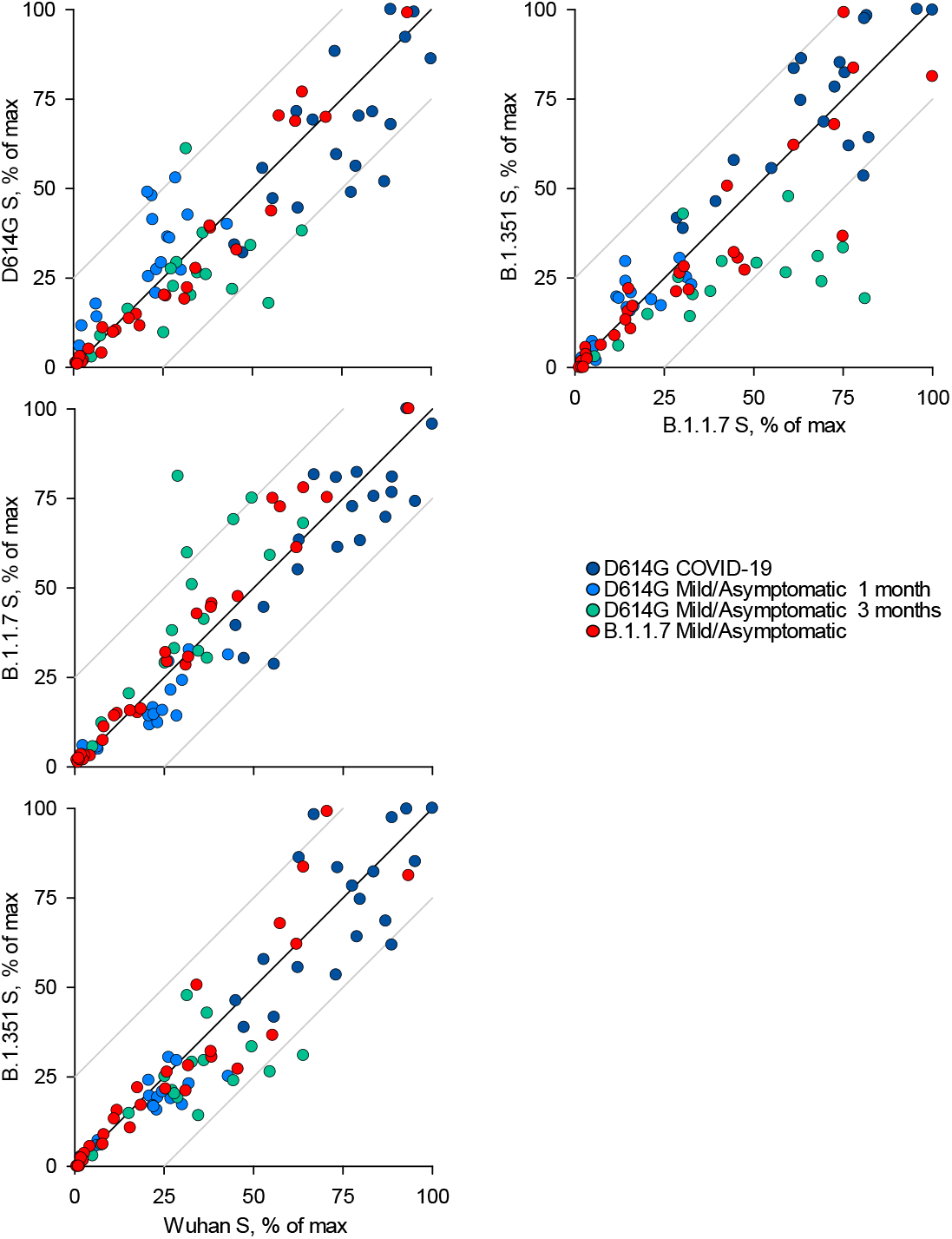
Recognition of distinct SARS-CoV-2 spike glycoproteins by antibodies in D614G and B.1.1.7 sera. Correlation of IgG antibody levels to Wuhan, D614G, B.1.1.7 and B.1.351 spikes in the indicated groups of donors infected either with the D614G or B.1.1.7 strains. Each symbol represents an individual sample and levels are expressed as a percentage of the positive control. Black lines denote complete correlation and grey lines a 25% change in either direction.

**Figure S3.**
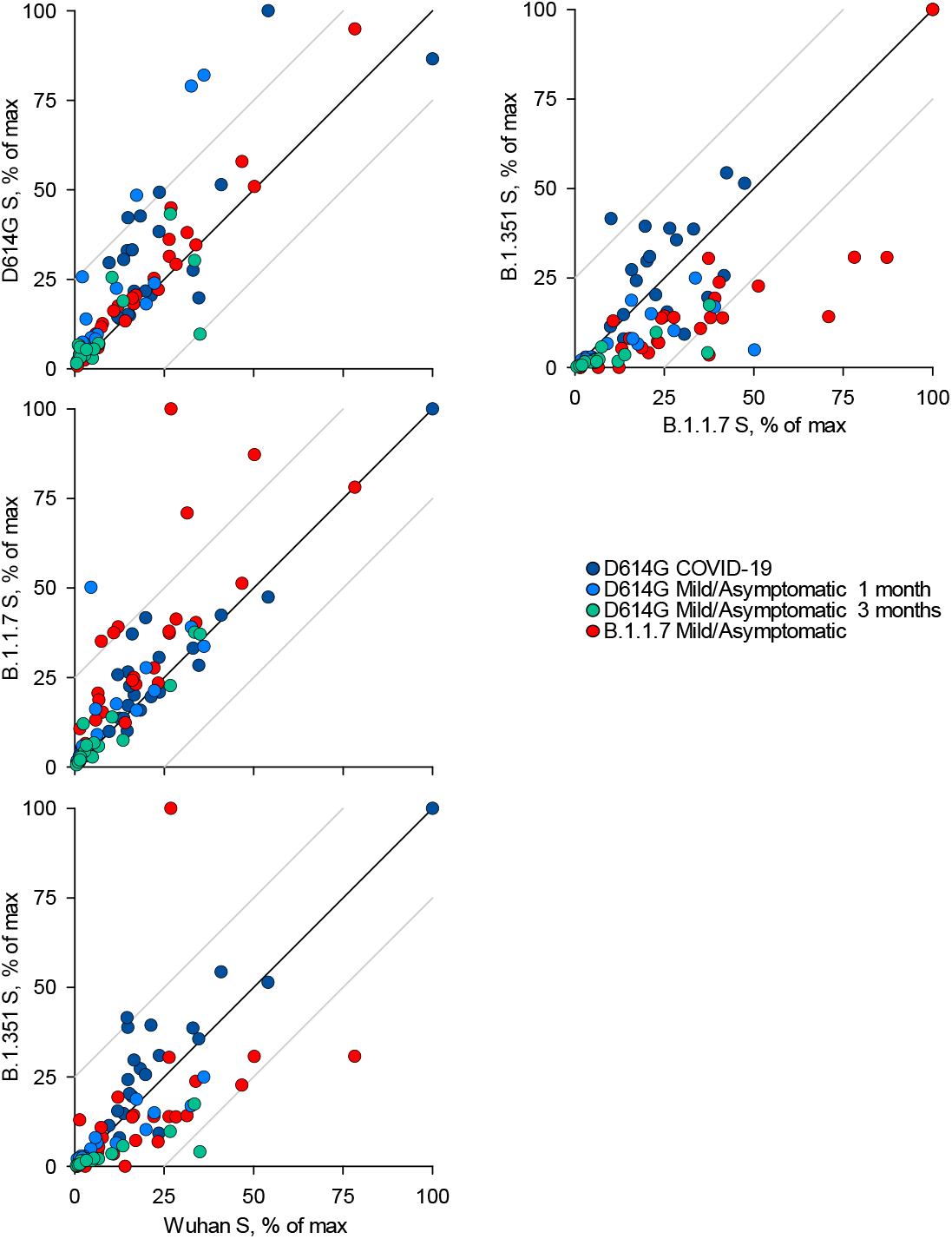
Recognition of distinct SARS-CoV-2 spike glycoproteins by antibodies in D614G and B.1.1.7 sera. Correlation of IgM antibody levels to Wuhan, D614G, B.1.1.7 and B.1.351 spikes in the indicated groups of donors infected either with the D614G or B.1.1.7 strains. Each symbol represents an individual sample and levels are expressed as a percentage of the positive control. Black lines denote complete correlation and grey lines a 25% change in either direction.

**Figure S4.**
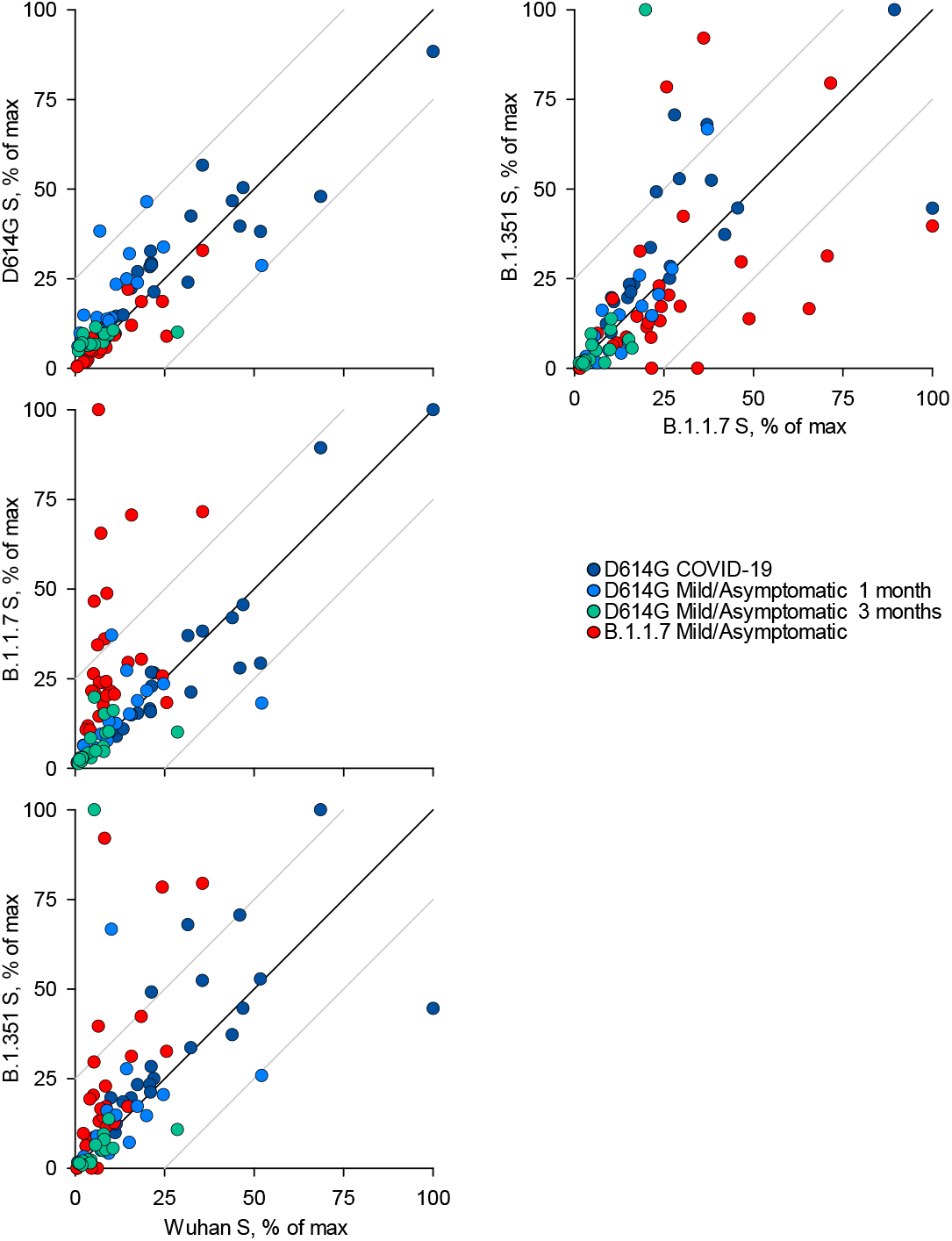
Recognition of distinct SARS-CoV-2 spike glycoproteins by antibodies in D614G and B.1.1.7 sera. Correlation of IgA antibody levels to Wuhan, D614G, B.1.1.7 and B.1.351 spikes in the indicated groups of donors infected either with the D614G or B.1.1.7 strains. Each symbol represents an individual sample and levels are expressed as a percentage of the positive control. Black lines denote complete correlation and grey lines a 25% change in either direction.

**Figure S5.**
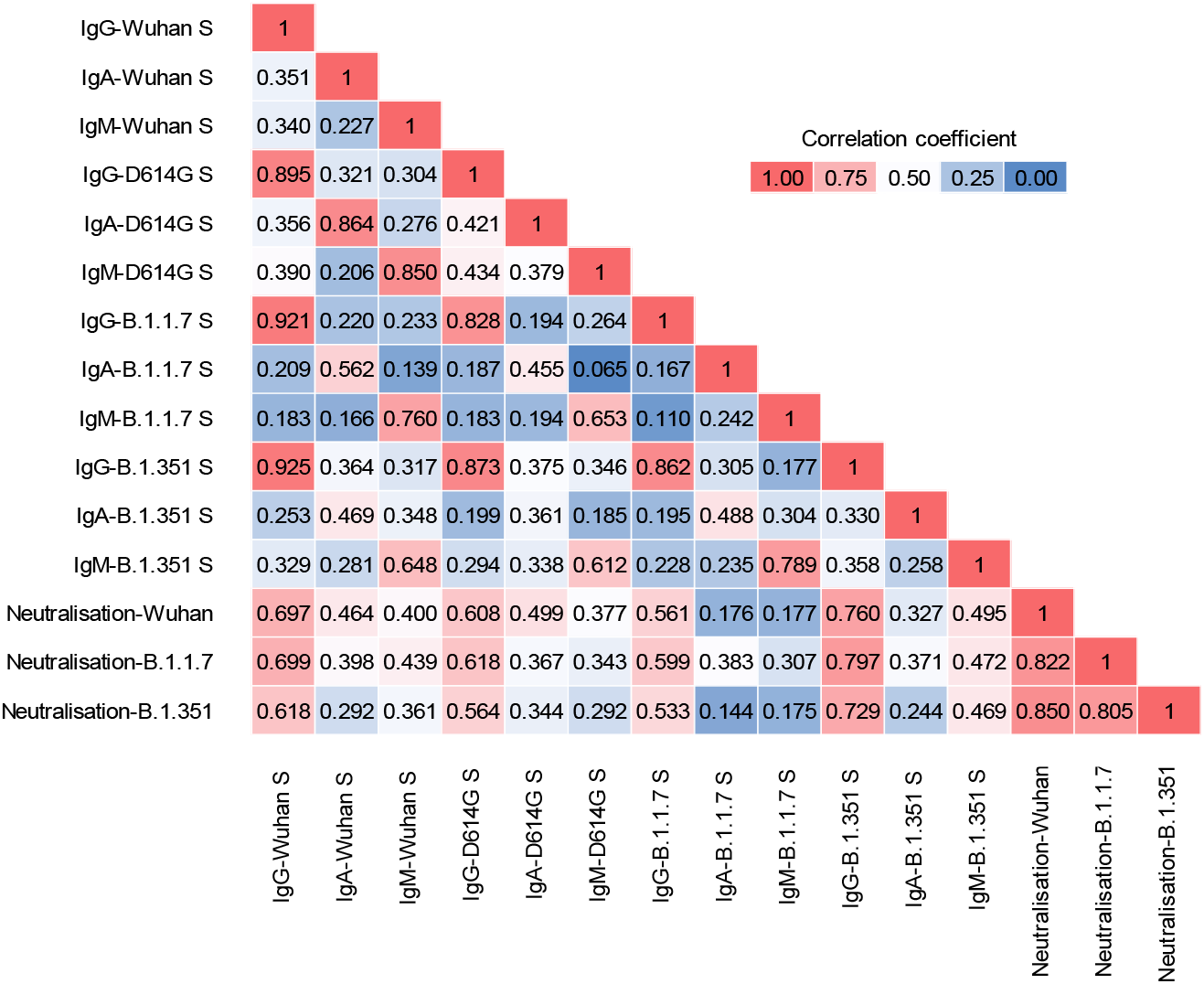
Matrix of correlation coefficients between binding and neutralising antibodies. Levels of binding IgG, IgM and IgA antibodies to the indicated spikes and levels of neutralising antibodies to the indicated strains were correlated using all the samples described this work (n=83).

**Figure S6.**
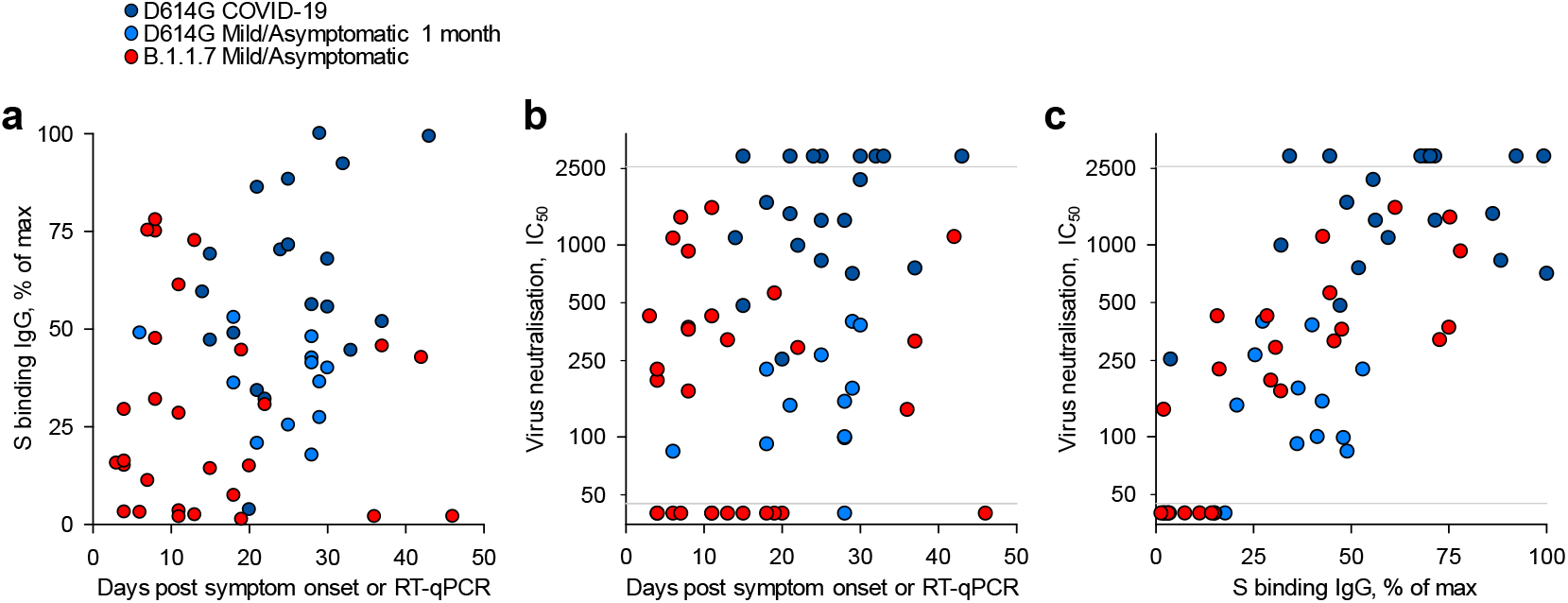
Kinetics and magnitude of the antibody response to D614G and B.1.1.7 infection. **a**, Levels of IgG antibodies to the spike of the infecting strain in sera from donors infected with the D614G or B.1.1.7 strains, over time since onset of symptoms (for symptomatic cases) or the first positive RT-qPCR diagnosis (for asymptomatic cases). Levels are expressed as a percentage of the positive control. **b**, Neutralising antibody levels (IC_50_) against the closest infecting strain (Wuhan for D614G infection) and B.1.1.7 for B.1.1.7 infection) in sera from donors infected with the D614G or B.1.1.7 strains, over time since onset of symptoms or since the first positive RT-qPCR diagnosis. **c**, Correlation of binding IgG and neutralising antibody levels from a and b, respectively. Grey horizontal lines denote the lower and upper limit of detection. In a-c, each symbol represents an individual sample.

**Figure S7.**
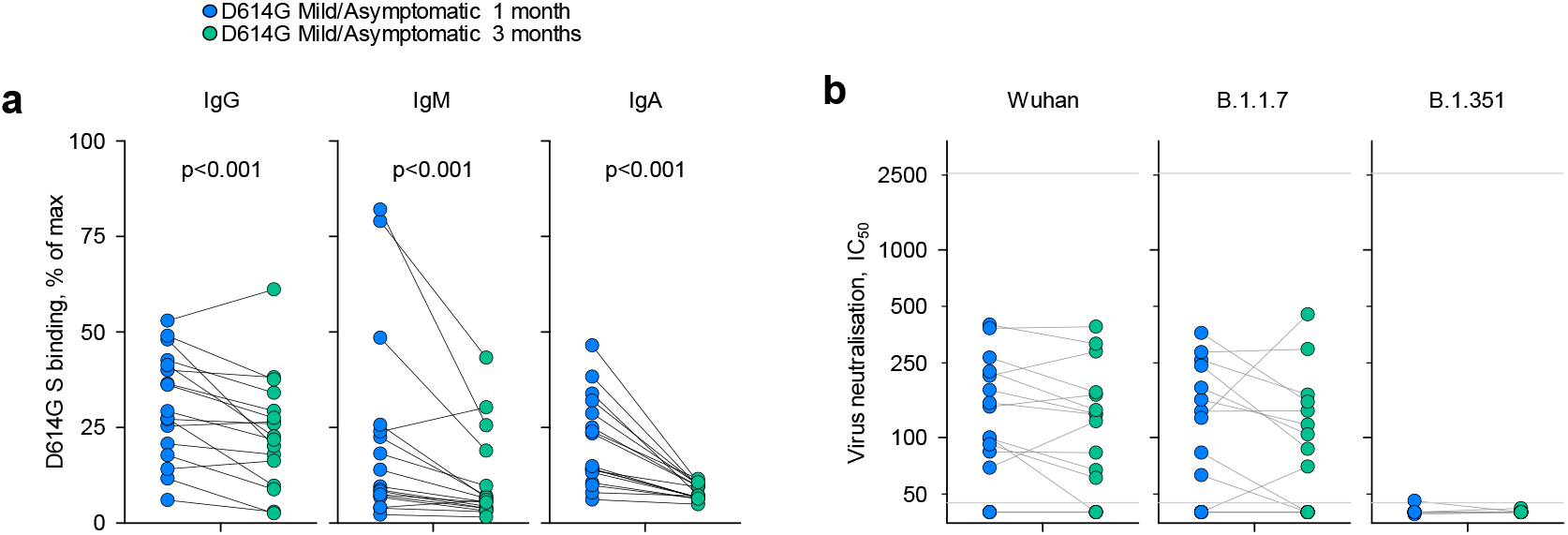
Persistence of binding and neutralising antibodies at a three-month follow-up of mild/asymptomatic D614G infection. **a**, Levels of IgG, IgM and IgA antibodies (expressed as a percentage of the positive control) to the D614G spike in sera from D614G-infected donors at one and three months post infection. **b**, Neutralising antibody levels (IC_50_) against the Wuhan, B.1.1.7 or B.1.351 strains in same donors describe in a. In a and b, connected symbols represent individual donors.

**Figure S8.**
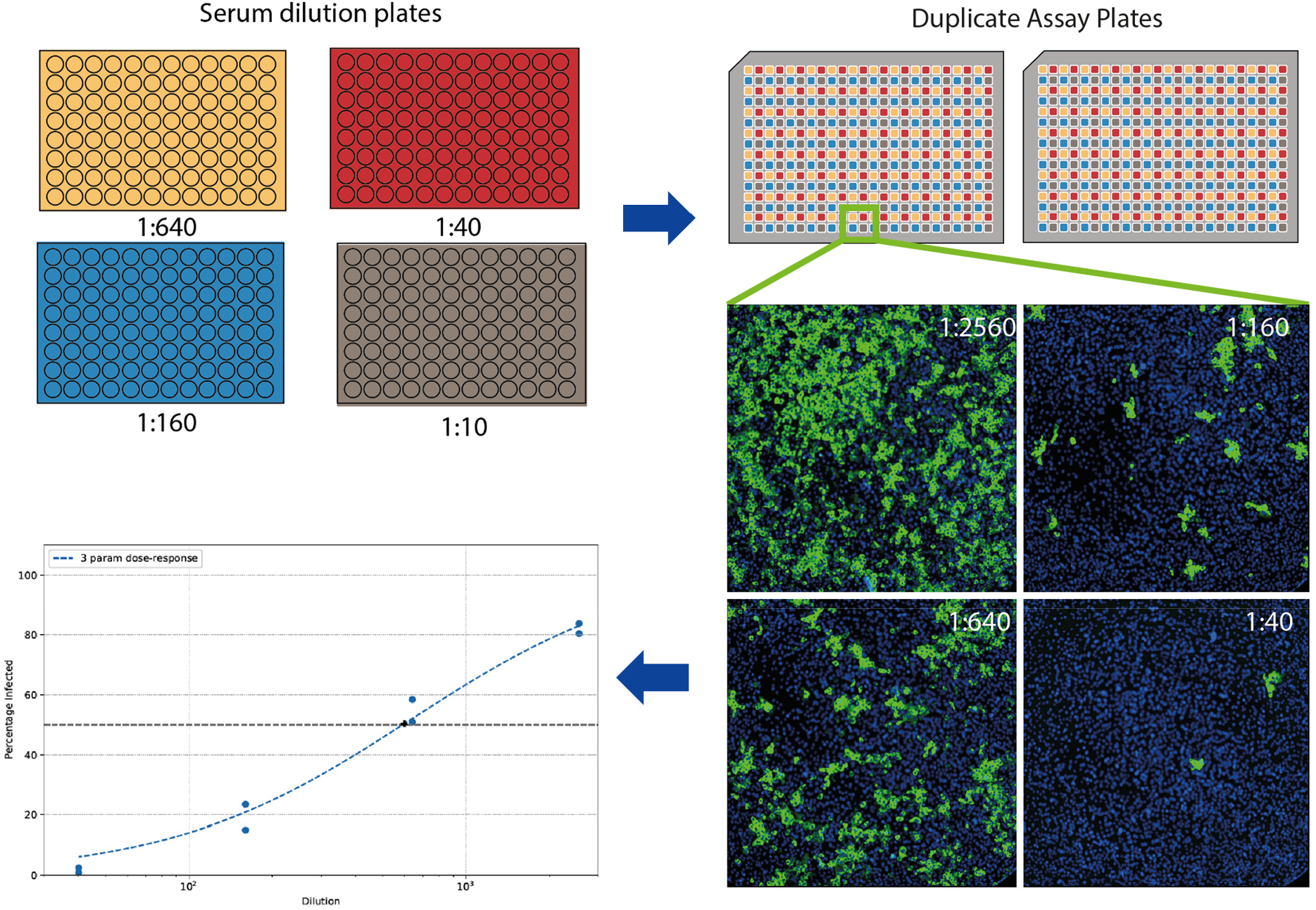
SARS-CoV-2 neutralisation assay set up. 96-well racks of serum samples including controls are serially diluted after an initial dilution of 1:10 to generate 4 total dilution plates. These are used to treat pre-seeded Vero E6 cells in 384-well assay plates in duplicate before infection with SARS-CoV-2 virus. After immunostaining with DAPI and a 488-conjugated monoclonal antibody against SARS-CoV-2 nucleoprotein, each well is imaged and infection area per area of cells calculated, followed by automated curve-fitting and identification of serum dilution factor to achieve 50% neutralisation (IC_50_).

